# Antimicrobial solid media for screening non-sterile *Arabidopsis thaliana* seeds

**DOI:** 10.1101/852020

**Authors:** James B. Y. H. Behrendorff, Guillem Borràs-Gas, Mathias Pribil

## Abstract

**Background:** Stable genetic transformation of plants is a low-efficiency process, and identification of positive transformants usually relies on screening for expression of a co-transformed marker gene. Often this involves germinating seeds on solid media containing a selection reagent. Germination on solid media requires surface sterilization of seeds and careful aseptic technique to prevent microbial contamination, but surface sterilization techniques are time consuming and can cause seed mortality if not performed carefully. We developed an antimicrobial cocktail that can be added to solid media to inhibit bacterial and fungal growth without impairing germination, allowing us to bypass the surface sterilization step.

**Results:** Adding a combination of terbinafine (1 µM) and timentin (200 mg/L) to solid media delayed the onset of observable microbial growth and did not affect germination of non-sterile seeds from ten different wild-type and mutant *Arabidopsis thaliana* accessions. The method was also compatible with *Nicotiana tabacum* germination. Seedlings sown in non-sterile conditions could be maintained on antimicrobial media for up to a week without observable contamination. The antimicrobial cocktail was compatible with rapid screening methods for hygromycin B, phosphinothricin (BASTA) and nourseothricin resistance genes, meaning that positive transformants can be identified from non-sterile seeds in as little as four days after stratification and transferred to soil before the onset of visible microbial contamination.

**Conclusion:** The antimicrobial cocktail presented here delays microbial growth for long enough to permit germination of non-sterile *Arabidopsis thaliana* seedlings on solid media and it is compatible with rapid screening methods. We were able to select genetic transformants on solid media without seed surface sterilization, eliminating a tedious and time-consuming step.

## Background

Driven by cheap and accessible methods of DNA assembly, the synthetic biology revolution has made it possible for molecular biologists to design and build dozens of new plasmids in as little as one or two weeks even without automation equipment. Plant science has not fully exploited these advances in molecular cloning to the same extent as other disciplines, partly because of experimental throughput limitations unique to plants.

Agrobacterium-mediated genetic transformation is one of the most versatile and accessible methods for modifying the genome of *Arabidopsis thaliana* [1, 2], but this approach produces only a small minority of seeds in the T_1_ generation that carry the transgene of interest. Transformation efficiencies between 0.57-2.57 % have been reported with optimized variations of the classic floral dipping method [2–4]. Identifying this minority of positive transformants usually relies on selection or screening for a co-transformed marker gene.

The most common selection approaches involve germinating seeds on an agar-based solid nutrient medium that contains a chemical reagent to select seedlings that express the corresponding marker gene. Popular selectable markers confer resistance to phosphinothricin (BASTA, also known as glufosinate), kanamycin, hygromycin B, or nourseothricin (also known as streptothricin) [5, 6]. Germination on solid media requires that seeds are surface sterilized to prevent overgrowth by microbial contaminants during the selection process.

At the time when these screening methods were established, molecular cloning was a bottleneck in the experimental workflow of transgenic plant preparation and typically few transgenic lines were prepared simultaneously. This is no longer the case, yet the same screening methods are still widely used. Screening for successful stable transfection events now represents a significant bottleneck, especially when an experiment involves several different genetic designs.

The seed sterilization step in particular has disadvantages that become more pronounced when screening increasing numbers of transformant lines. Liquid sterilization in hypochlorite bleach has a low seed mortality rate but is tedious, requiring several washing steps that become time consuming when preparing large quantities of seeds [7]. Chlorine gas is suitable for sterilizing seeds from multiple lines simultaneously, but gas sterilization still requires up to four hours of waiting time and can have a relatively high mortality rate even when the gas concentration is carefully controlled [7]. Mortality caused by the sterilization process could result in the loss of rare transformants or a reduction in the diversity of mutant libraries. Furthermore, surface sterilization does not necessarily eliminate microbial spores that can be trapped inside the seed coat during embryogenesis [8].

We aimed to develop a method that would allow us to avoid surface sterilization of seeds altogether. Our approach was to identify a combination of antifungal and antibacterial compounds that inhibit microbial growth but do not impair *Arabidopsis* germination and growth. When combining the method presented here with established rapid selection methods [5], we were able to identify positive transformants from non-sterile seeds and transfer them to soil for propagation prior to the onset of observable microbial contamination.

## Results

### Terbinafine as an antifungal reagent

Terbinafine is an antifungal reagent that inhibits squalene epoxidase, causing a deficiency in the membrane lipid ergosterol [9]. Squalene epoxidation is also a key step in the biosynthesis of plant sterols, and squalene epoxidase knockout mutants of *A. thaliana* exhibit increased sensitivity towards terbinafine [10]. We sought to test whether low concentrations of terbinafine could be used to inhibit fungal growth without impairing germination of *A. thaliana*.

In a preliminary experiment, non-sterile seeds from four wild-type *A. thaliana* ecotypes (Columbia (Col-0), Landsberg *erecta* (Ler-0), Wassilewskija (Ws-0), and Nossen (No-0)), six photosynthetic gene mutants (*curt1abcd* [11], *atpC1* [12], *hcf136* [13], *pam68* [14], *psaL* [15], and *npq4* [16]), and *Nicotiana tabacum* (cv. Petit Havana) were sown directly onto 0.5X Murashige and Skoog (MS) agar with sucrose (1 %, w/v) and terbinafine (added to a final concentration of 1, 0.1, or 0.01 μM). Negative control plates contained DMSO (0.1 %, v/v) without terbinafine. While sucrose is not necessary for germination of wild-type plants, many mutants with impaired photosynthesis benefit from the addition of sucrose during germination. Sucrose also increases the risk of microbial contamination because it is a utilizable carbon source for most fungi and many bacteria. Therefore, we included sucrose in our media to ensure that our antimicrobial medium would be useful in cases where the inclusion of sucrose is necessary.

Seeds were stratified by wrapping the agar plates with aluminium foil and storing them at 4 °C for 68 h. The foil was removed after stratification and plates were transferred to growth chambers. The onset of germination (defined as the first cotyledons to emerge on each plate) was determined by visual inspection and was scored qualitatively, as was the emergence of observable microbial contamination. Plates were inspected twice per day for seven days (168 h in total).

Terbinafine did not affect the onset of germination at any of the concentrations tested (up to 1 μM) (Additional File 1: Supporting Figure S1A). In negative control agar plates that lacked terbinafine, microbial contamination was observed as early as 24 h after being transferred to growth chamber conditions (median time to visible contamination: 64 h) (Additional File 1: Supporting Figure S1B). In the presence of terbinafine, there was a general trend toward delayed onset of microbial contamination with increasing terbinafine concentration. At 1 µM terbinafine, all plates were free of microbial contamination after 168 h except for one plate, where a contaminant emerged after 112 h.

### Adding antibacterial β-lactam antibiotics

β-lactam antibiotics were tested as the antibacterial reagent because they inhibit peptidoglycan biosynthesis in prokaryotes, whereas most other classes of prokaryote-targeting antibiotics also interfere with plastid [17, 18] and mitochondrial protein synthesis [19]. We examined the effects of carbenicillin and timentin, which are both commonly used for eliminating Agrobacteria from plant tissue culture [2, 20, 21]. Timentin is a mixture containing a β-lactam antibiotic (ticarcillin) and a β-lactamase inhibitor (clavulanic acid).

Timentin (200 mg/L) or carbenicillin (500 mg/L) was added to 0.5X MS agar that contained sucrose (1 %, w/v) and terbinafine (1 µM). Non-sterile seeds were sown directly onto agar plates and stratified as described above, and then transferred to growth chambers and monitored by visual inspection. Germination was quantified by recording the number of germinated seedlings twice per day for the first four days, and once per day thereafter. Microbial contamination was recorded qualitatively.

The combination of terbinafine (1 µM) and timentin (200 mg/L) did not inhibit germination of wild-type *A. thaliana* ecotypes or *N. tabacum* compared with untreated seeds (sown on 0.5X MS agar + sucrose without antimicrobial additives) (Fig. 1). Photosynthetic mutant *A. thaliana* lines were also unaffected (Fig. 2) except in the case of the *psaL* mutant, where the inclusion of timentin may have impaired germination in 7 % of seeds (at 168 h: 96 % of untreated seeds had germinated versus 89 % of seeds sown on timentin plus terbinafine). Carbenicillin (500 mg/L) inhibited normal root development in all lines examined and delayed germination in all cases except for the *curt1abcd* quadruple mutant, which exhibits slower germination than the wild-type ecotypes examined here and naturally produces a lower proportion of viable seeds [22]. The combination of terbinafine (1 µM) and timentin (200 mg/L) prevented microbial contamination for five days in 100 % of cases, and for seven days in 90 % of cases (Fig. 3). Henceforth we describe 0.5X MS agar containing this combination of terbinafine and timentin as MSTT agar, or MSTT+suc agar when the medium also contains sucrose (1 % w/v).

**Figure 1.**
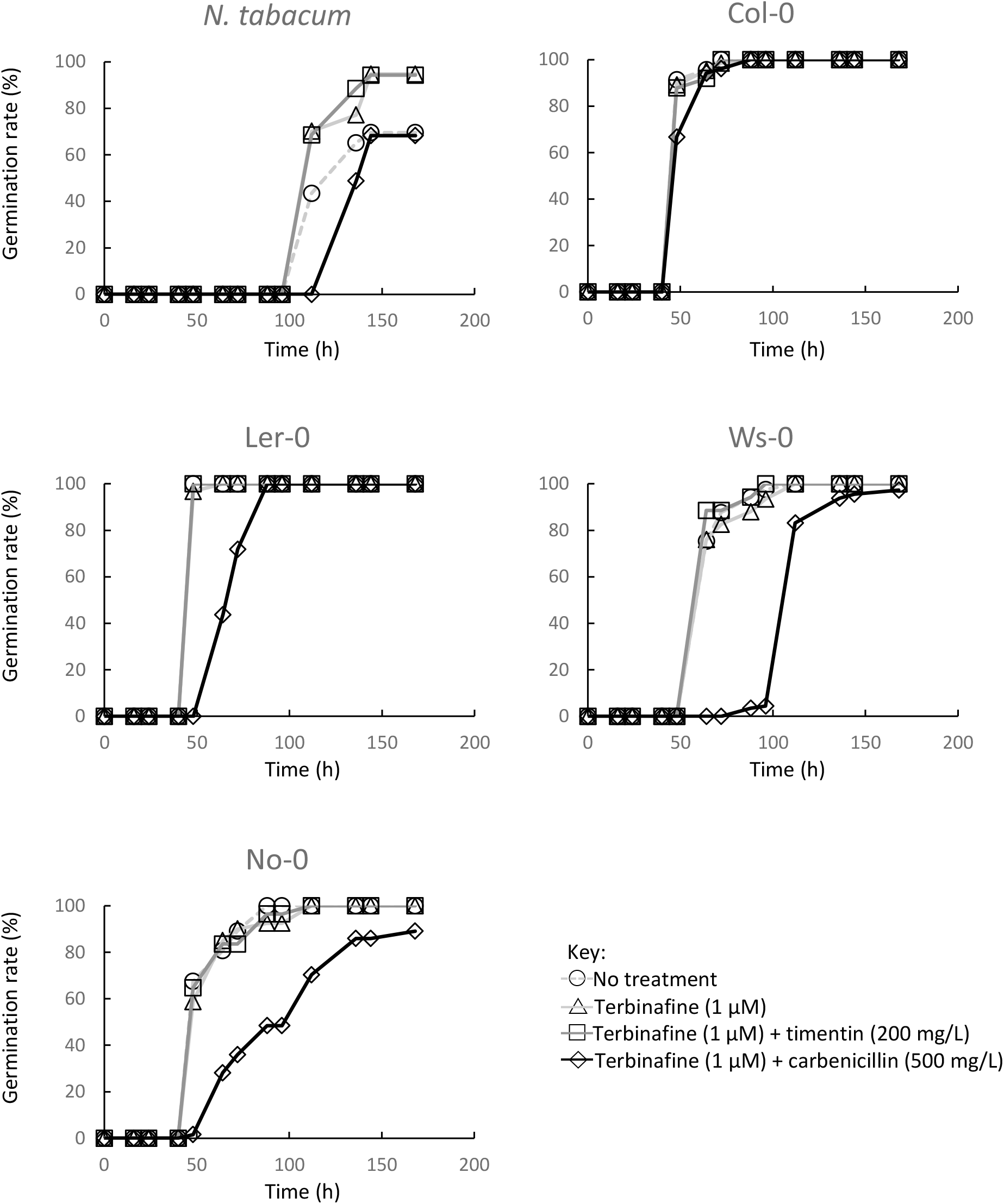
Germination of wild-type seeds in the presence of terbinafine and β-lactam antibiotics. Non-sterile seeds for *Nicotiana tabacum* (cv. Petit Havana) and *Arabidopsis thaliana* ecotypes Columbia (Col-0), Landsberg *erecta* (Ler-0), Wassilewskija (Ws-0), and Nossen (No-0) were sown on 0.5X MS agar with added sucrose (1 %, w/v) and different combinations of terbinafine and timentin or carbenicillin (indicated). Germination was monitored by visual inspection and the number of germinated seeds was recorded as a percentage of the total seeds sown on that agar plate.

**Figure 2.**
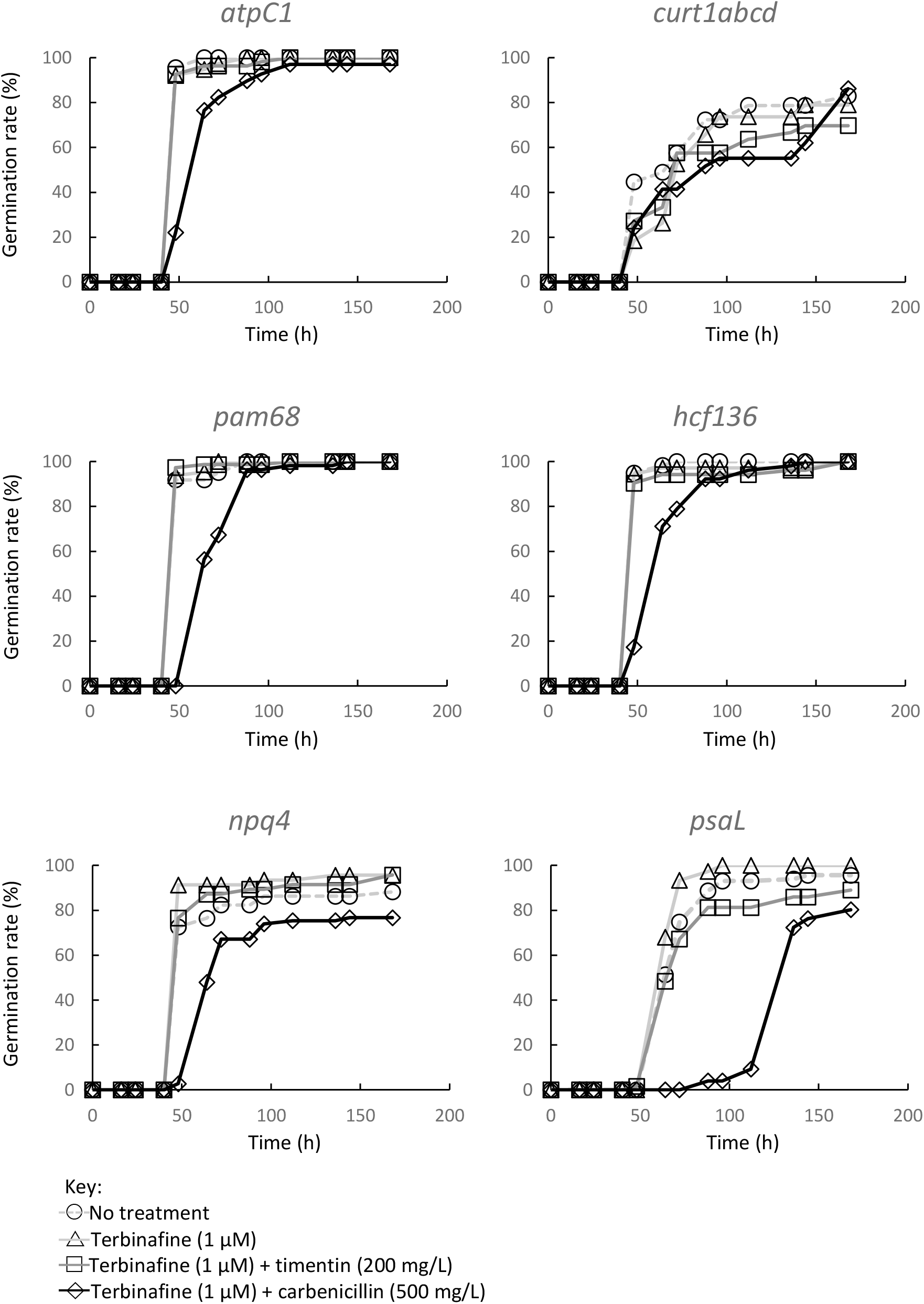
Germination of *A. thaliana* photosynthetic mutants in the presence of terbinafine and β-lactam antibiotics. Non-sterile seeds for six *A. thaliana* photosynthetic mutants (*atpC1*, *curt1abcd*, *pam86*, *hcf136*, *npq4*, and *psaL*) were sown on 0.5X MS agar with added sucrose (1 %, w/v) and different combinations of terbinafine and timentin or carbenicillin (indicated). Germination was monitored by visual inspection and the number of germinated seeds was recorded as a percentage of the total seeds sown on that agar plate.

**Figure 3.**
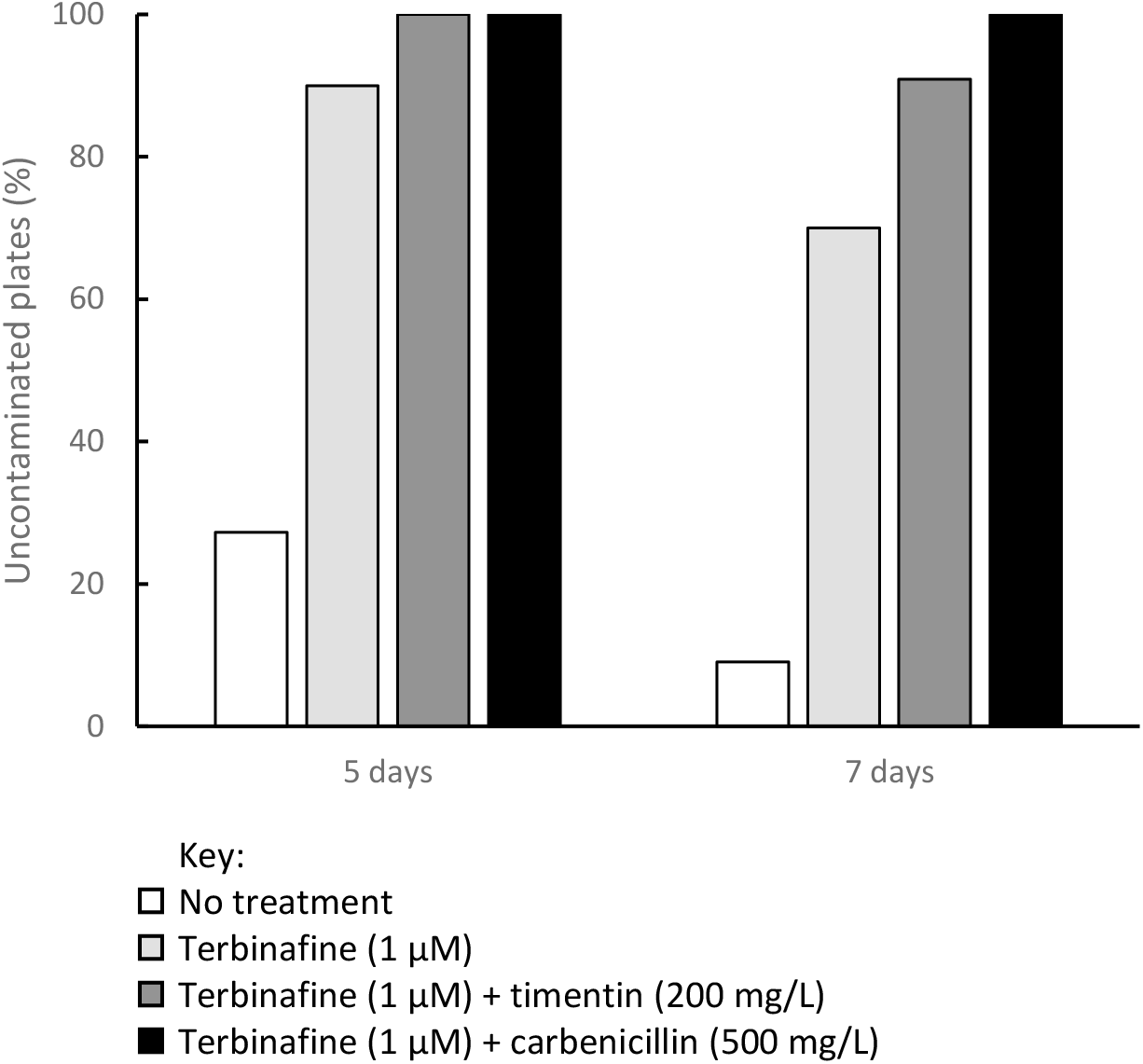
Onset of microbial contamination in the presence of terbinafine and β-lactam antibiotics. Non-sterile seeds were sown on 0.5X MS agar with added sucrose (1 %, w/v) and different combinations of terbinafine and timentin or carbenicillin (indicated). The proportion of agar plates that remained uncontaminated after five and seven days was recorded (n = eleven agar plates per condition).

Germination of the four wild-type *A. thaliana* ecotypes and *N. tabacum* was also examined on MSTT agar without sucrose, and in all cases germination was unaffected (Additional File 1: Supporting Figure S2). Contamination emerged on negative control plates (0.5X MS agar) after only 48 h, whereas MSTT agar plates remained free of observable contamination for one week (microbial contamination emerged on all MSTT agar plates after 184 h).

### Non-sterile screening for *Arabidopsis* transformants on selective agar

#### Screening for nourseothricin resistance and fluorescent protein expression

As a test case to see whether MSTT agar is useful for screening genetic transformants without seed sterilisation, we transfected *A. thaliana* (Col-0) with a green fluorescent protein expression construct (pN_35S/mEGFP: a monomeric enhanced green fluorescent protein (mEGFP) under the control of the cauliflower mosaic virus 35S promoter, with a nourseothricin acetyl transferase (*nat*) selectable marker) (Fig. 4A). Non-sterile seeds collected from the T_0_ plant were sown directly onto MSTT agar with added nourseothricin (50 mg/L). Seeds were stratified directly on the agar plates and then screened using the rapid hypocotyl elongation method that was previously developed for identifying hygromycin B resistance [5]. Briefly, this method involves exposing stratified seeds to light for 6 h to break dormancy, and then keeping the germinating seedlings in darkness at 22 °C to promote hypocotyl elongation.

**Figure 4.**
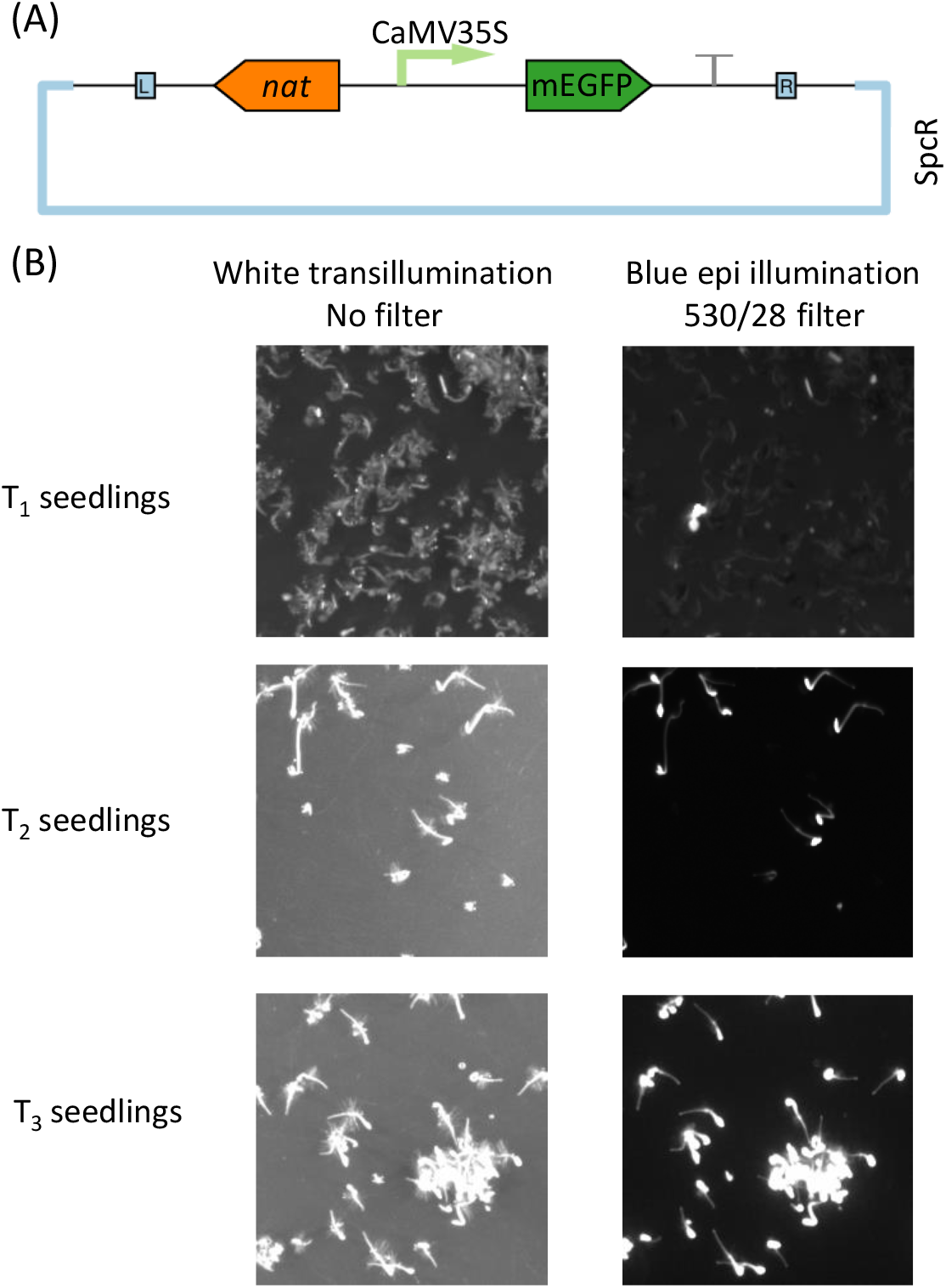
Screening for nourseothricin-resistant transformants with non-sterile seeds. *A. thaliana* (Col-0) was transfected with a green fluorescent protein expression construct and the resulting seeds were screened for transgene integration in non-sterile conditions on MSTT agar with added nourseothricin (50 mg/mL). (A) Schematic map of the plasmid used for Agrobacterium-mediated transfection. The T-DNA region is flanked by L and R, indicating the left and right border sequences. Monomeric enhanced green fluorescent protein (mEGFP) expression is regulated by the cauliflower mosaic virus 35S promoter (CaMV35S). Nourseothricin acetyl transferase (*nat*) is the selectable marker for plant transformation, and SpcR indicates that the plasmid backbone confers resistance to spectinomycin in bacteria. (B) The screening procedure identified positive transformant T_1_ plants and was repeated to identify homozygous plants in the T_3_ generation. Transgene integration and expression was confirmed by screening for hypocotyl elongation (indicating *nat* expression) and mEGFP fluorescence (imaged by illumination with a blue light source and a 530 nm filter with a 28 nm bandpass).

Plates were uncovered after two full days of dark treatment (i.e. on the fourth day post-stratification). Individual seedlings with extended hypocotyls were clearly identifiable amongst the majority of seedlings that did not have extended hypocotyls, and fluorescence imaging revealed mEGFP expression (Fig. 4B) in the same seedlings that had extended hypocotyls. No microbial growth was observed and several positive transformants were transferred to soil for propagation. This demonstrated that the hypocotyl elongation method can be used to screen for nourseothricin resistance, and that nourseothricin-based screening is compatible with the use of terbinafine and timentin to limit microbial growth. Homozygous transformant lines were identified by repeating the screening procedure with seeds collected from T_1_ and T_2_ plants (Fig. 4B).

We used the same method to identify plants that overexpress the mApple red fluorescent protein (transformed with pN_35S/mApple) and a nuclear-encoded chloroplast-targeted mCitrine yellow fluorescent protein (pN_35S/CTP-mCitrine) (Additional File 1: Supporting Figure S3).

#### Screening for BASTA resistance and fluorescent protein expression

We also tested whether non-sterile germination on MSTT agar could be coupled with rapid phosphinothricin (BASTA)-based screening. *A. thaliana* (Col-0) plants were transfected with the pB_35S/mEGFP expression construct (identical to pN_35S/mEGFP except that it has a phosphinothricin N-acetyltransferase (*bar*) selectable marker in place of the nourseothricin acetyl transferase marker) (Fig. 5A). Seeds from the T_0_ plant were sown directly onto MSTT agar with added BASTA (50 µM) and stratified as described above. BASTA-resistant seedlings were identified using the previously-published rapid screening method [5] with minor modifications. Briefly, stratified seeds were exposed to light for 6h to break dormancy, then they were kept in darkness at 22 °C for three days before transferring to long day growth chamber conditions (see *Methods* for details). After two days in growth chamber conditions, transformed seedlings were clearly identifiable by their dark green expanded cotyledons while non-transformed seedlings exhibited pale unexpanded cotyledons. These phenotypic differences became more pronounced after three days. Fluorescence imaging confirmed mEGFP expression in seedlings with green expanded cotyledons (Fig. 5B). No microbial growth was observed during the screening period and multiple positive transformants were identified.

**Figure 5.**
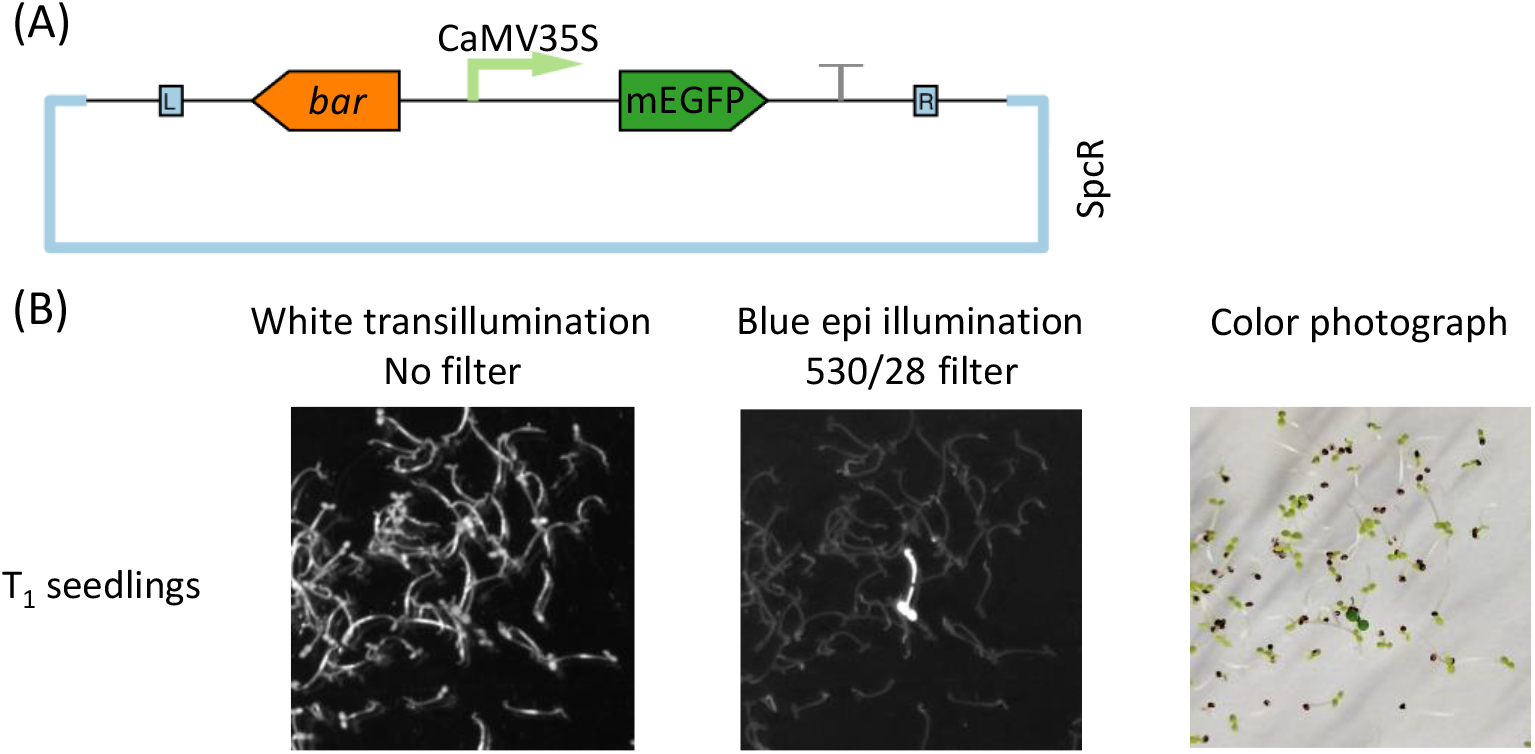
Screening for BASTA-resistant transformants with non-sterile seeds. *A. thaliana* (Col-0) was transfected with a green fluorescent protein expression construct and the resulting seeds were screened for transgene integration in non-sterile conditions on MSTT agar with added BASTA (50 μM). (A) Schematic map of the plasmid used for Agrobacterium-mediated transfection. The T-DNA region is flanked by L and R, indicating the left and right border sequences. Monomeric enhanced green fluorescent protein (mEGFP) expression is regulated by the cauliflower mosaic virus 35S promoter (CaMV35S). Phosphinothricin *N*-acetyl transferase (*bar*) is the selectable marker for plant transformation, and SpcR indicates that the plasmid backbone confers resistance to spectinomycin in bacteria. (B) The screening procedure identified positive transformant T_1_ plants. Transgene integration and expression was confirmed by screening for cotyledon expansion and greening (indicating *bar* expression) and mEGFP fluorescence (imaged by illumination with a blue light source and a 530 nm filter with a 28 nm bandpass).

#### Screening for hygromycin B resistance and identifying mutants produced with CRISPR-Cas9

In a third test case, we transfected *A. thaliana* (Col-0) with a construct for CRISPR-Cas9-mediated functional knockout of the phytoene desaturase (*PDS3*) gene, which causes an albino phenotype [23]. Our plasmid (GS2.1/EC) combined a previously-published *PDS3*-targeting sgRNA [24] with a Cas9 gene under the control of a previously-published egg cell-specific promoter [25]. The T-DNA region also carried a hygromycin phosphotranferase gene conferring resistance to hygromycin B (Fig. 6A).

**Figure 6.**
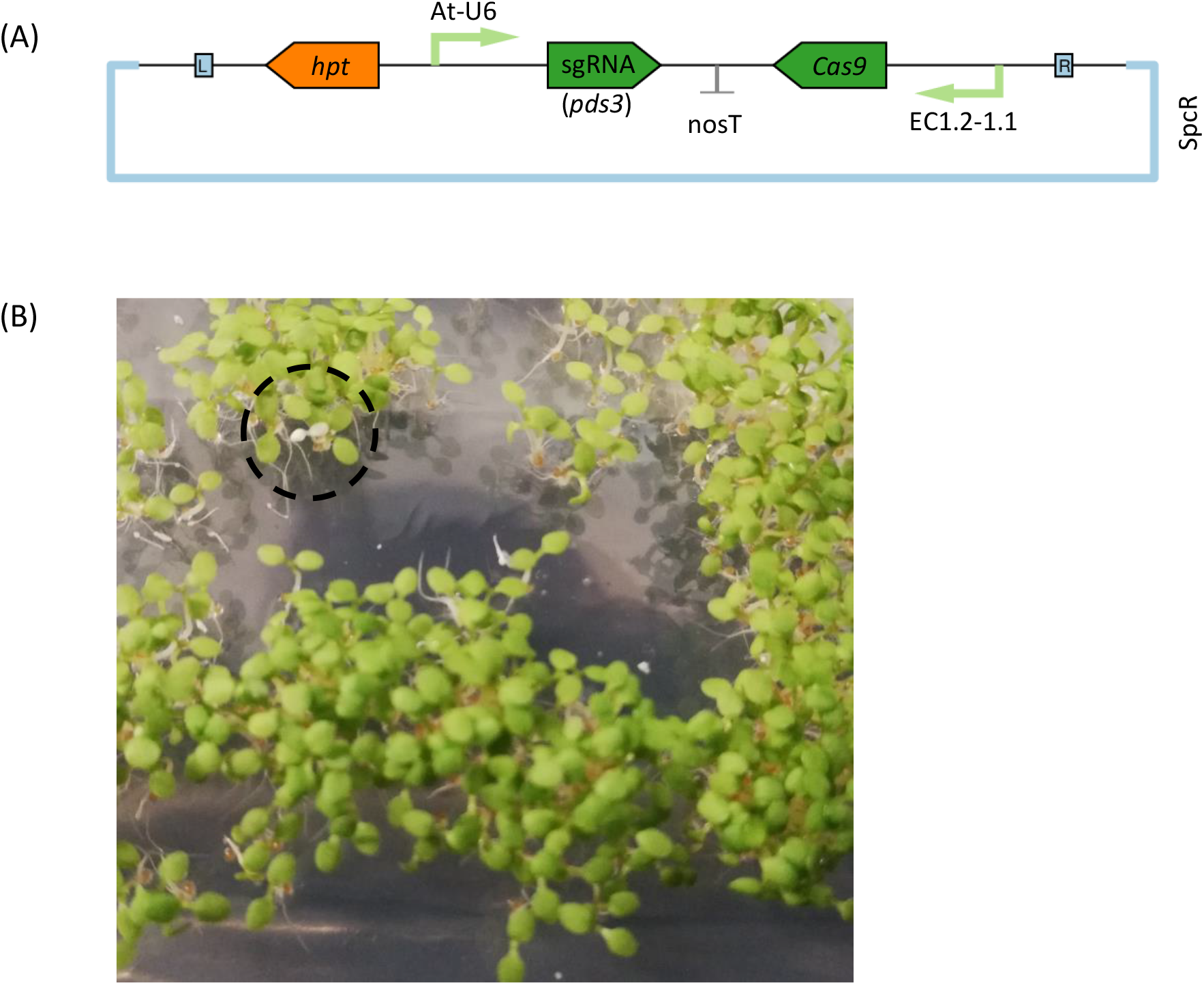
Screening for a CRISPR-Cas9-mediated *pds3* mutant phenotype with non-sterile seeds. *A. thaliana* (Col-0) was transfected with a CRISPR-Cas9 plasmid targeting mutation of the phytoene desaturase, *PDS3*. Homozygous *pds3* mutants were obtained by screening seeds in non-sterile conditions on MSTT+suc agar with added hygromycin B (15 mg/L). (A) Schematic map of the plasmid used for Agrobacterium-mediated transfection. The T-DNA region is flanked by L and R, indicating the left and right border sequences. A single guide RNA (sgRNA) targeting *PDS3* is regulated by the *A. thaliana* U6 polymerase III promoter (At-U6). The Cas9 gene is regulated by an egg cell-specific promoter (EC1.2-1.1). The hygromycin phosphotransferase (*hpt*) is the selectable marker for plant transformation, and SpcR indicates that the plasmid backbone confers resistance to spectinomycin. (B) Non-sterile seeds were screened on MSTT+suc agar with hygromycin B. Homozygous *pds3* knockout mutants were identifiable in the T_2_ generation by their characteristic albino phenotype. An example *pds3* mutant on day five post-stratification is indicated inside the dashed circle.

Non-sterile seeds collected from T_0_ plants were sown on MSTT+suc agar with hygromycin B (15 mg/L). Positive transformants were identified on the basis of hypocotyl elongation on the fourth day post-stratification. T-DNA integration was confirmed by PCR analysis of leaf tissue (Additional File 1: Supporting Figure S4), but all positive transformants in the T_1_ generation had green cotyledons, indicating that any post-transfection CRISPR-Cas9 activity during seed development had not resulted in homozygous *pds3* mutants.

Non-sterile seeds collected from one T_1_ plant were sown on MSTT+suc agar containing hygromycin B (15 mg/L). Approximately 1500 seeds were sown on a single 12 cm x 12 cm agar plate. Approximately 75 % of seedlings were resistant to hygromycin B, and on the fifth day post-stratification (i.e. after cotyledon greening) three albino *pds3* mutants were identified (Fig. 6B). The three albino seedlings were confirmed as independent *pds3* mutants by Sanger sequencing (Additional File 1: Supporitng Figure S5).

## Discussion

When producing new *A. thaliana* transgenic lines, experimental throughput is partly limited by the transformant screening process. In particular, the seed sterilization step is time consuming and can cause seed mortality [7]. Alternative screening methods that negate the need for seed sterilization have been developed, each with advantages and disadvantages.

Conventional selection for BASTA resistance (conferred by *bar*, the phosphinothricin N-acetlytransferase gene) [26] involves spraying the aerial parts of germinated seedlings and can be performed in non-sterile conditions with seeds sown directly on soil at relatively high densities. The disadvantage of this approach is the time required to identify positive transformants: typically at least three spray applications are spread across three weeks [27]. Additionally, the use of BASTA is restricted in some countries due to neurotoxicity linked to BASTA ingestion [28–30].

A more modern approach to avoiding seed sterilization is to use a fluorescent protein with seed-specific expression as the marker gene [31, 32]. Accumulation of the fluorescent protein in transformed seeds can be observed visually with a suitable light source and filter combination (a fluorescence microscope is typically used). Visual screening for a co-expressed fluorescent protein avoids the need for sterilization, and only seeds with active transgene expression are sown on soil. This is an excellent approach for avoiding seed mortality and reducing the number of plants that need to be grown in a screening campaign, but this approach is still best suited to scenarios involving relatively few transgenic lines due to the labour involved in screening seeds under a fluorescence microscope.

The method we present here provides another pragmatic option for avoiding seed sterilization. When paired with rapid screening methods, seeds sown on MSTT or MSTT+suc agar could be screened for resistance to nourseothricin, hygromycin B or BASTA in as few as 4-5 days after stratification. This approach allowed us to curate homozygous T_3_ transgenic lines and identify CRISPR-Cas9-mediated *pds3* knockout mutants without the need for seed sterilization at any stage. It is possible that the selection reagents contribute to the antimicrobial effect when used in combination with terbinafine and timentin, but nourseothricin and hygromycin B were not sufficient to prevent contamination when used alone (data not shown).

Although MSTT and MSTT+suc agar did not negatively affect germination of any of the seeds examined in this study, we only validated these media for screening laboratory-grown seeds for transgene insertion. It is unknown whether exposure to sub-inhibitory concentrations of terbinafine and timentin may trigger any responses in *Arabidopsis* that would make these media unsuitable for physiological studies. Additionally, all seeds used in this study were grown in growth chamber conditions; our method may not be suitable for use with field-grown seeds that could be expected to have a greater microbial burden.

While the inhibitory effect of terbinafine on squalene epoxidase in plants has been characterised, the effects of β-lactam antibiotics in plants are not fully understood. Peptidoglycan biosynthesis is retained in moss chloroplasts but is absent from vascular plants [33], and it is generally assumed that β-lactam antibiotics do not affect the chloroplasts of higher plants [34]. Carbenicillin (500 mg/L) has been described in the literature as beneficial for eliminating β-lactam-sensitive *Agrobacterium* strains from transfected *Arabidopsis* and tobacco tissue culture [20], and carbencillin concentrations between 100-500 mg/L have also been recommended for use in solid media when screening T_1_ transgenic *Arabidopsis* seeds after floral dip transformation[2, 21, 35]. We initially planned to use a high concentration of carbenicillin (500 mg/L) on the basis that β-lactamase enzymes are secreted by many environmental bacteria and some commonly used laboratory strains of Agrobacteria [36]. However, carbenicillin and penicillin were recently reported to impair root elongation in *A. thaliana* at concentrations between 100-1000 mg/L [37]. As an alternative to carbenicillin, we considered timentin on the basis that a lower concentration of timentin should provide a similar protective effect due to the presence of a β-lactamase inhibitor (clavulanic acid) in the timentin formulation. We observed that carbenicillin (500 mg/L) did not cause seed mortality but delayed germination and prevented root elongation, whereas timentin (200 mg/L) had no observable effect on *Arabidopsis* germination and growth.

We believe that our method provides another useful approach to simplifying *Arabidopsis* transformant screening. It requires minimal labour and seeds can been sown at high densities when using hypocotyl elongation-based rapid screening for nourseothricin or hygromycin B resistance. It can be used with rapid screening methods for BASTA (and potentially kanamycin) resistance [5] if seeds are sown at sufficiently low density to distinguish between green (positive transformant) and pale yellow (wild-type) cotyledons.

## Conclusions

Timentin and terbinafine added to 0.5X MS agar delay the onset of microbial contamination and do not inhibit germination of *A. thaliana* or *N. tabacum*. The inhibition of microbial growth is sufficient to allow selection of transgenic plants from non-sterile seeds, avoiding the time-consuming seed sterilization step and minimizing seed mortality.

## Methods

### Chemicals

Murashige and Skoog medium including vitamins (MS medium) [38] (Cat. no. M0222), hygromycin B (Cat. no. H0192) and timentin (ticarcillin 2NA and clavulanate K 15:1 mixture, Cat. no. T0190) were purchased from Duchefa Biochemie. Nourseothricin was purchased from Jena Bioscience (Cat. no. AB-102L). BASTA was purchased from Bayer Cropscience (Product no. 84442615). Terbinafine (Cat. no. T8826) was purchased from Merck. All other chemicals were the highest quality locally available. Stock solutions were prepared as follows: timentin, 200 mg/mL in water; nourseothricin, 50 mg/mL in water; terbinafine, 1 mM in dimethylsulfoxide (DMSO); carbenicillin, 50 mg/mL in water; BASTA, 50 mM in water. Hygromycin B was used directly from the liquid stock provided by the manufacturer (400 mg/mL in water).

### Plant accessions

*Arabidopsis thaliana* mutants (*curt1abcd* [11], *atpC1* [12], *hcf136* [13], *pam68* [14], *psal* [15], and *npq4* [16]) and wild-type ecotypes (Columbia (Col-0), Landsberg *erecta* (Ler-0), Wassilewskija (Ws-0), and Nossen (No-0)) were a gift from Prof. Dario Leister, Ludwig Maximilians Universität, Germany. *Nicotiana tabacum* (cv. Petit Havana) seeds were a gift from Dr Lars Scharff (University of Copenhagen, Denmark).

### Antimicrobial compound screening

0.5X MS agar was prepared by dissolving Murashige and Skoog medium in water and adjusting the pH to 5.7 with 1 M potassium hydroxide. Agar was added to a final concentration of 1 % (w/v) and sterilized by autoclaving. Sucrose was added from a filter-sterilized stock solution (40 % w/v in water). Antimicrobial reagents were added from the stock solutions described above (in *Chemicals*), and all agar plates were poured in non-sterile conditions on a standard laboratory bench. Non-sterile seeds were sown directly onto solidified agar and stratified by wrapping the agar plates in aluminium foil and storing at 4 °C for 68 h. After stratification, all agar plates were maintained under long day conditions in a growth chamber (16 h light at 100 μmol m^-2^s^-1^, 8 h darkness, 22 °C, 60 % humidity).

The final antimicrobial solid medium developed in this study (MSTT agar) contained 0.5X MS medium, terbinafine (1 μM), timentin (200 mg/L), and agar (10 g/L). MSTT+suc agar also contained sucrose (1 %, w/v).

### Plasmids

All cloning strategies were designed with Geneious 10.2.6 (http://www.geneious.com) and performed using the general principles of the Gibson assembly method [39]. Cartoon representations of plasmids were generated with Pigeon [40] (http://pigeon.synbiotools.org). Oligonucleotide primers and sources of template DNA are listed in Additional File 2. *Fluorescent reporter protein constructs*: in a previous study [41], a nourseothricin-selectable plasmid was prepared by replacing the bialaphos resistance (*bar*) gene from plasmid pB2GW7 [42] with the nourseothricin acetyl transferase (*nat*) gene from *Streptomyces noursei*. The resulting plasmid is described as pN_35S. The ccdB counter-selectable marker was replaced with the coding sequences for either mEGFP or mApple fluorescent proteins, placing fluorescent protein expression under the control of the cauliflower mosaic virus 35S promoter (plasmids pN_35S/mEGFP and pN_35S/mApple available at www.addgene.org as plasmids #132565 (RRID:Addgene_132565) and #132566 (RRID:Addgene_132566), respectively). The pB_35S/mEGFP plasmid was prepared by cloning the mEGFP coding sequence directly into the pB2GW7 backbone, replacing the ccdB counterselectable marker (www.addgene.org, plasmid #135320 (RRID:Addgene_135320)). We have also made available a hygromycin-selectable mEGFP plasmid, pH_35S/mEGFP (www.addgene.org, plasmid #135321 (RRID:Addgene_135321)), prepared by cloning the mEGFP coding sequence into the pH2GW7 [42] backbone. Plasmid pN_35S/CTP-mCitrine was produced in an earlier study [41] and encodes an mCitrine fluorescent protein fused in-frame to the chloroplast transit peptide from RuBisCO small subunit 1A (www.addgene.org, plasmid #117989 (RRID:Addgene_117989)). *CRISPR constructs*: an mApple fluorescent protein was fused in-frame to the C-terminus of a *Streptomyces pyogenes* Cas9 via a GGGGS flexible linker. The Cas9-mApple coding sequence and a *PDS3* sgRNA under the control of the *Arabidopsis thaliana* U6 polymerase III promoter [24] were cloned into the pH2GW7 [42] backbone (hygromycin selection). A previously-described promoter made by combining two *A. thaliana* egg cell specific promoters [25] was then inserted upstream of the Cas9-mApple coding sequence to create GS2.1/EC (www.addgene.org, plasmid #132568 (RRID:Addgene_132568)).

Plasmids were transformed into *Agrobacterium fabrum* strain GV3101 (previously known as *Agrobacterium tumefaciens* GV3101 [43]) via electroporation with the following conditions: voltage 2500 V, capacitance 25 μF, resistance 400 Ω, 2 mm cuvette.

### Genetic transformation and screening

*A. thaliana* (Col-0) was grown under long day conditions (16 h light at 100 μmol m^-2^s^-1^, 8 h darkness) at 22 °C and 60 % humidity and transformed according to a modified floral dip method described previously [4]. Seeds collected from transformed plants were sown on MSTT or MSTT+suc agar with appropriate selection reagents.

Plants resistant to nourseothricin (50 mg/L) or hygromycin B (15 mg/L) were identified by rapid screening for hypocotyl elongation [5]. Seeds were stratified directly on agar plates, which were wrapped in aluminium foil and stored at 4 °C for 68 h. After stratification, plates were shifted to growth chamber conditions and exposed to light for six hours. Plates were then wrapped in foil to maintain darkness for two full days and stored at 22 °C. On the fourth day, plates were unwrapped and resistant seedlings with elongated hypocotyls were clearly distinguishable from non-resistant seedlings. Plants resistant to phosphinothricin (50 µM) were identified by rapid screening for green expanded cotyledons [5]. Plates were shifted to growth chamber conditions after stratification and exposed to light for six hours, and then wrapped in foil to maintain darkness for three full days. Following the dark treatment, plates were unwrapped and kept in a growth chamber under long day conditions. Positive transformants could be identified two days later (i.e. the fifth day post-stratification), and differences between resistant and non-resistant seedlings were more pronounced after three days.

In the case of plants transformed with fluorescent protein expression constructs (pN_35S/mEGFP, pN_35S/CTP-mCitrine, pN_35S/mApple, or pB_35S/mEGFP), transformation was verified by fluorescence imaging on a Bio-Rad ChemiDoc XRS+ (Bio-Rad Laboratories, Inc.). Green and yellow fluorescent signals (from mEGFP and chloroplast-targeted mCitrine) were captured using blue light epi-illumination and a 530 nm filter (28 nm bandpass). Red fluorescence from mApple was captured with green light epi illumination and a 605 nm filter (50 nm bandpass). For red fluorescence imaging, it was necessary to image seedlings on the fourth day post-stratification prior to greening of cotyledons. It was not possible to identify mApple fluorescence in green leaves due to interference from chlorophyll autofluorescence.

Homozygous *pds3* mutants were identified by their distinct albino phenotype and were confirmed via Sanger sequencing (oligonucleotide primer details included in Additional File 2).

### PCR from leaf tissue

Diagnostic PCRs, preparative PCRs for Sanger sequencing, and PCRs to prepare *A. thaliana* DNA for cloning were performed using leaf tissue as the source of template DNA. A portion of leaf tissue (approximately 5 mm^2^) was homogenized by grinding in 50 μL of 1X Q5 reaction buffer (New England Biolabs Cat. No. B9027S) in a 1.5 mL microcentrifuge tube with a micropestle. The homogenate was heated to 98 °C for 10 min, then cooled on ice. After cooling, leaf debris was separated by centrifugation (30 s, 13,000 *g*). The supernatant was used directly as a source of template DNA (1 μL template DNA per PCR).

## Declarations

### Ethics approval and consent to participate

Not applicable.

### Consent for publication

All authors have read the final version of the manuscript and approve its submission for publication.

### Availability of data and material

Plasmids created for this study are available at www.addgene.org using the reference numbers described in the text and summarised in Additional File 2. Raw data are available at the following URL: https://www.dropbox.com/s/l9ze4qxcthgq0wo/Behrendorff%20et%20al%20antimicrobials%20raw%20data.zip?dl=0.

### Authors’ contributions

JBYHB conceived of the concept, designed and executed the experiments, and wrote the manuscript. GBG validated the use of MSTT agar for BASTA-based screening. MP contributed to experimental design and writing the manuscript.

## Acknowledgements

We thank Dr Omar Sandoval-Ibañez (Max Planck Institute of Molecular Plant Physiology, Germany) and Dr Lars Scharff (University of Copenhagen, Denmark) for their feedback on the manuscript.

