## Additional File 1 for "Antimicrobial solid media for screening non-sterile *Arabidopsis thaliana* seeds"

Fig. S1

(A)

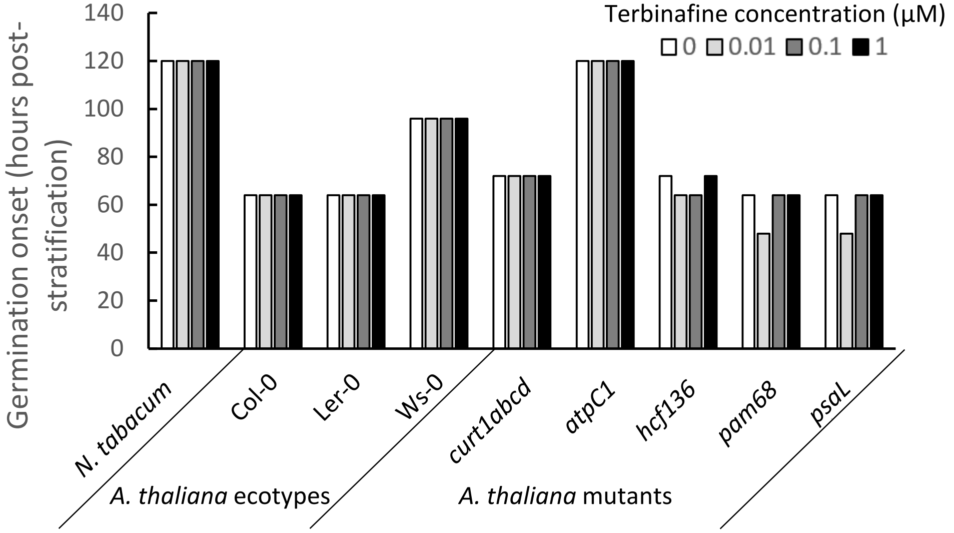


(B)

| Terbinafine concentration (µM) | 0 | 0.01 | 0.1 | 1 |
|---|---|---|---|---|
| Contamination frequency (plates) | 9/9 | 8/9 | 6/9 | 1/9 |

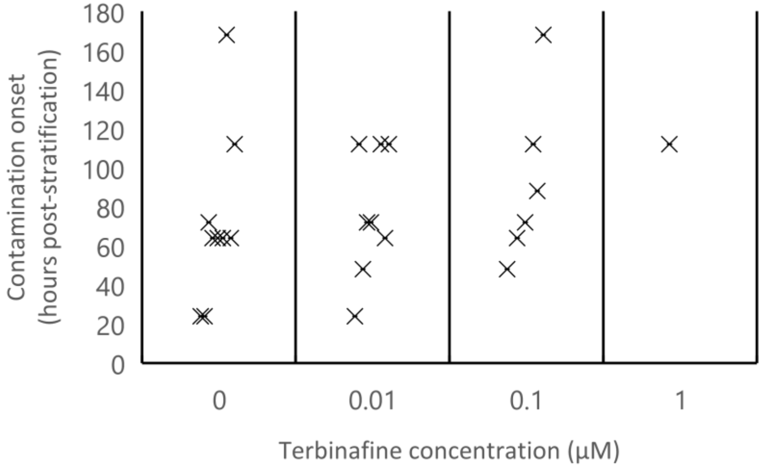


Supporting Figure S1 **Terbinafine as an antifungal reagent.** Non-sterile seeds were sown on 0.5X MS agar with added sucrose (1 %, w/v) and different concentrations of terbinafine (indicated). (A) Germination onset is defined as the emergence of cotyledons from the first germinating seeds on each agar plate. (B) Each marker (×) indicates the time at which microbial contamination emerged on individual agar plates. The overall frequency of contamination is summarised at the top of the plot.

Fig. S2

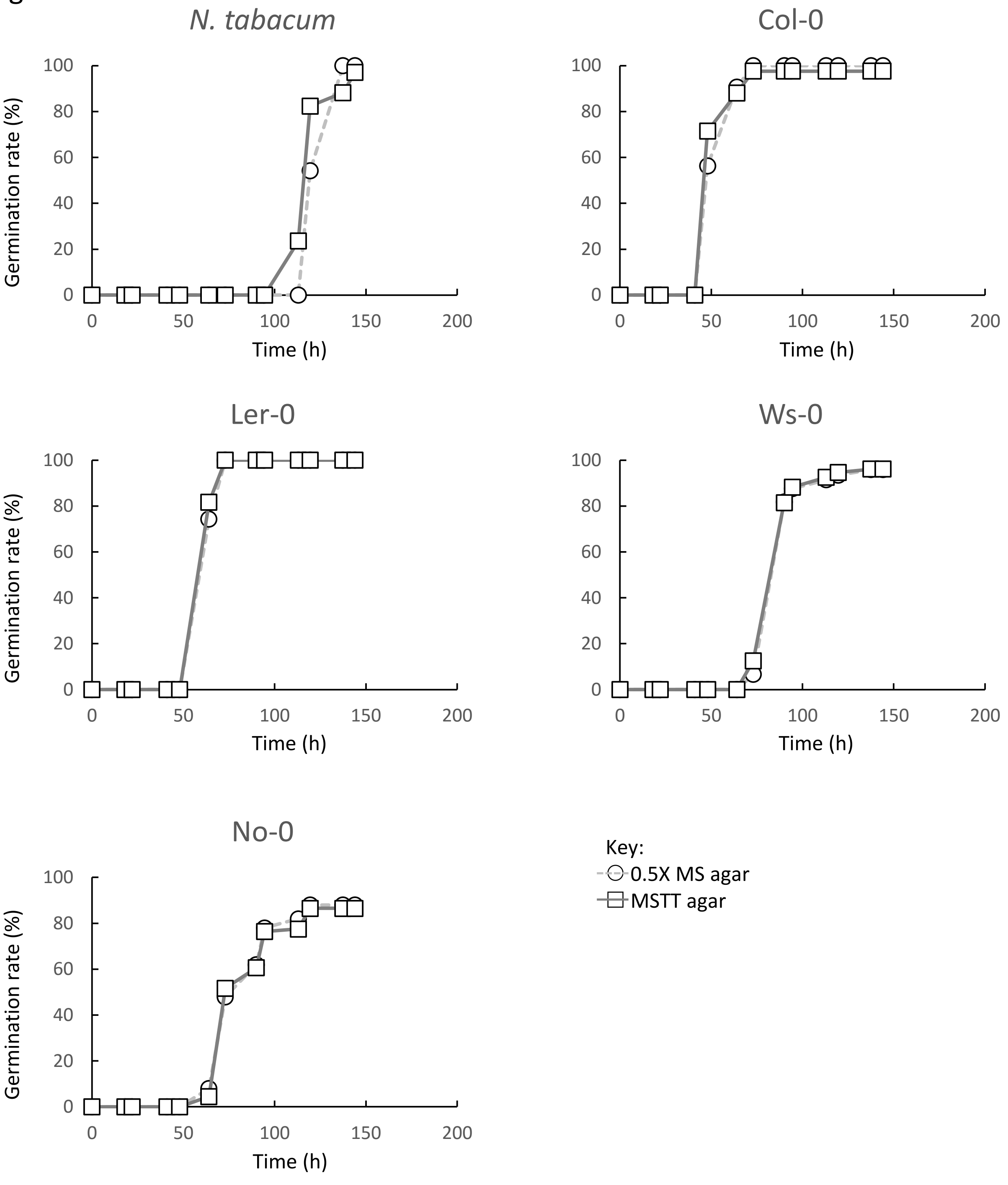


Supporting Figure S2 **Germination of non-sterile wild-type seeds on MSTT agar.** Non-sterile seeds of *N. tabacum* and four *A. thaliana* ecotypes were sown on 0.5X MS agar (○) or MSTT agar (□). Plates were observed twice per day and the proportion of germinated seeds was recorded. Germination was defined as cotyledon emergence.

Fig. S3

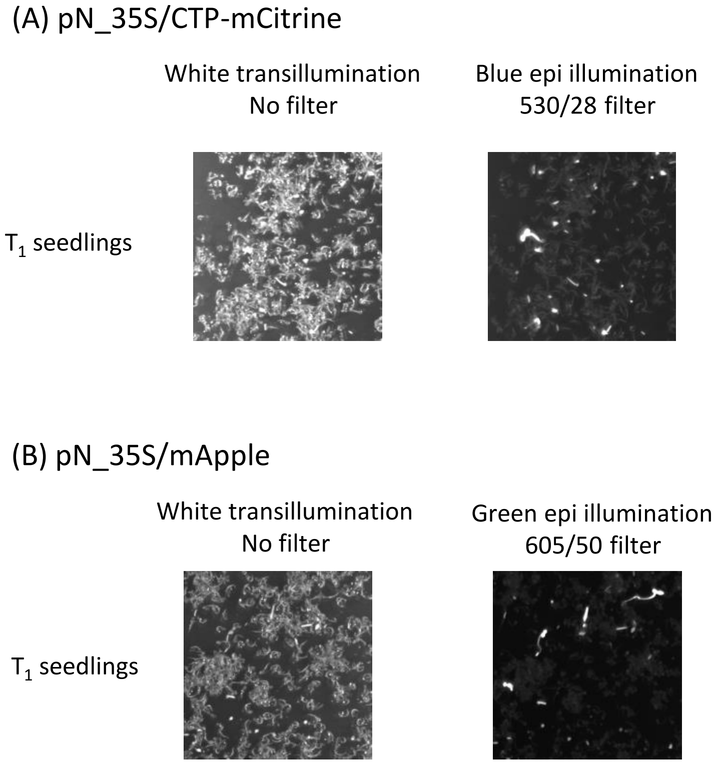


Supporting Figure S3 **Verification of transgene expression with fluorescence imaging.** Positive transformants for nuclear-encoded chloroplast-targeted mCitrine expression (A) and mApple expression (B) were verified with fluorescence imaging.

Fig. S4

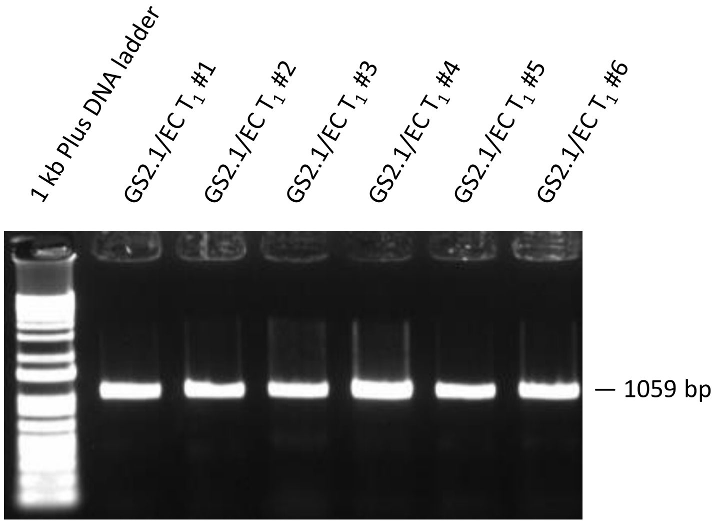


Supporting Figure S4 **Verification of Cas9 presence in positive GS2.1/EC transformant lines.** Six independent $T_1$ plants were identified that showed resistance to hygromycin B. Leaf tissue was sampled for PCR with primers specific to a 1059 bp section of the Cas9 gene. Agarose gel electrophoresis of the PCR products is shown. Gel layout: 1 kb Plus DNA ladder (ThermoFisher Scientific Cat. no. 10787018), $T_1$ plants 1-6.

Fig. S5

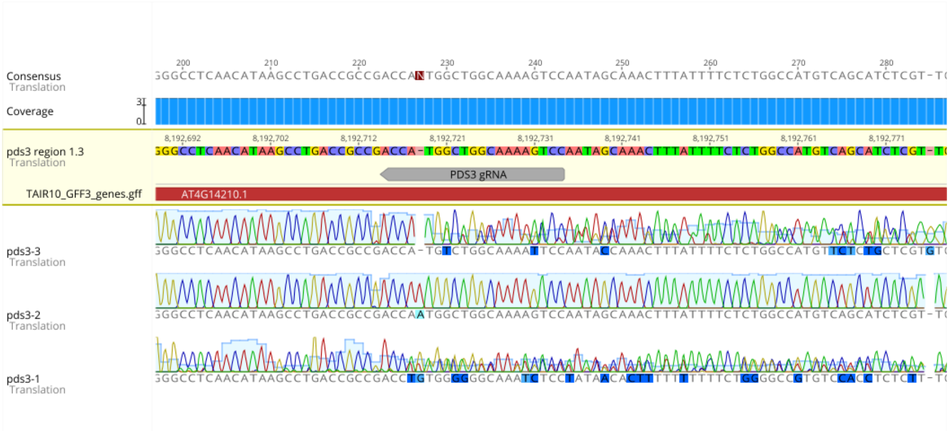


Supporting Figure S5 **Verification of *pds3* knockouts.** CRISPR-Cas9-mediated knockout of the *pds3* gene was performed by stable transfection of *A. thaliana* Col-0 with the GS2.1/EC construct. The targeted region of *pds3* from three albino *A. thaliana* mutants was sequenced. All three mutants are independent knockout lines. *pds3*-2 is homozygous for a single nucleotide insertion, while *pds3*-1 and *pds3*-3 appear to have heterozygous knockout mutations (i.e. different mutations on each chromosome), indicated by mixed chromatogram peaks after the apparent Cas9 cut site.
